# Sex-, strain and lateral differences in brain cytoarchitecture across a large mouse population

**DOI:** 10.1101/2022.08.09.503434

**Authors:** David Elkind, Hannah Hochgerner, Etay Aloni, Noam Shental, Amit Zeisel

## Abstract

The mouse brain is by far the most intensively studied among mammalian brains, yet basic measures of its cytoarchitecture remain obscure. For example, quantifying cell numbers, and the interplay of sex-, strain-, and individual variability in cell density and volume is out of reach for many regions. The Allen Mouse Brain Connectivity project produces high-resolution full brain images of hundreds of brains. Although these were created for a different purpose, they reveal details of neuroanatomy and cytoarchitecture. Here, we used this population to systematically characterize cell density and volume for each anatomical unit in the mouse brain. We developed a deep neural network-based segmentation pipeline that uses the auto-fluorescence intensities of images to segment cell nuclei even within the densest regions, such as the dentate gyrus. We applied our pipeline to 537 brains of males and females from C57BL/6J and FVB.CD1 strains. Globally, we found that increased overall brain volume does not result in uniform expansion across all regions. Moreover, region-specific density changes are often negatively correlated with the volume of the region, therefore cell count does not scale linearly with volume. Many regions, including layer 2/3 across several cortical areas, showed distinct lateral bias. We identified the greatest strain-specific or sex-specific differences in the medial amygdala (MEA), bed nuclei (BST), lateral septum and olfactory system (e.g., MOB, AOB, TR) and prefrontal areas (e.g., ORB) – yet, inter-individual variability was always greater than the effect size of a single qualifier. We provide the results of this analysis as an accessible resource for the community.

## Introduction

The mammalian brain can be divided into neuroanatomical units (i.e., brain regions) characterized by a shared function, connectivity, developmental origin, and/or cytoarchitecture (i.e., number and density of cells it contains). The mouse brain is the most extensively studied in mammals and its regions are well characterized. Although cytoarchitecture is one of the most prominent features of a brain region, few studies have systematically mapped cell bodies or quantified cell densities in mouse brains, compared to the early, detailed cell mapping of the nematode *C. elegans*.^1^

Obtaining an accurate cell count for a brain region is technically challenging. Previous estimates relied heavily on extrapolation from manual counting of 2D sections (stereology), making cell-resolved data for subcortical regions sparse ^2^. Analyzing complete brains using 2D histological sections remains labor-intensive because it requires sectioning, mounting, and accurate alignment with a reference atlas. Furthermore, automated cell counting proved particularly difficult in dense regions, such as the hippocampal formation and the cerebellum ^3^. Automated block-face imaging methods solved several of these issues and drastically improved throughput ^4^, For instance, serial two-photon tomography (STPT) ^5^ was a technological breakthrough that integrated tissue sectioning with top-view light microscopy. STPT provided high-quality imaging in an optical plane below the sectioning surface, and solved many problems of section distortion and atlas alignment, further easing downstream analysis. Yet, STPT typically represents a subsample of the complete volume, and some interpolation is needed.

Because of their limited throughput, histological studies cannot supply the number of analyzed brains needed to uncover potential variability between individuals, experimental conditions, and populations. Complementary approaches aimed at evaluating variability, e.g., magnetic resonance imaging (MRI), can measure some features, such as the volume of given brain regions, and can even track individuals over time in a noninvasive manner. Yet, MRI lacks the accuracy needed for counting cells or assessing cell densities, and it remains difficult to simultaneously analyze regional volume *and* cell density with high accuracy brain-wide, especially in the large throughput required for comparing two experimental populations (such as two strains, or males vs. females). Therefore, there is a need for systematic measurement of all cells over hundreds of brains from multiple experimental groups.

To address this knowledge gap, we harnessed the Allen Mouse Brain Connectivity Project (AMBCP) dataset, which is largest existing cohort of whole-brain STPT images, produced by the Allen Institute for the different purpose of mapping mouse regional connectivity ^6^. We applied a deep neural network (DNN) to discern cell nuclei, using their background autofluorescence channel. This enabled us to perform a systematic brain-wide cell density estimation across hundreds of mouse brains. Based on the alignment with the Allen Mouse Brain Atlas (AMBA), we were able to simultaneously measure volume *and* density for each brain, for each region, over a large population. We constructed a comprehensive database that aggregates these results and provides them as an accessible resource to the community. We also discovered non-trivial relationships between densities and volumes, and gained insights into strain- and sex-dependent characteristics across various brain regions.

## Results

### Autofluorescence of STPT images displays cell nuclei

The AMBCP project was first published in 2014 ^6^. The project has systematically imaged 2,992 full brains, using serial two-photon tomography, for the purpose of tracing neuronal projections and mapping regional (mesoscale) connectivity with GFP-labelled viral tracers. Each brain in the dataset is covered by 130-140 (median 137) serial coronal sections, with a gap of 100μm, as reported in the AMBCP study (Allen Mouse Brain Connectivity Atlas Documentation, 2017). We noticed that the red (background) channel of STPT images, taken for the purpose of atlas alignment, typically features dark, round-like objects resembling cell nuclei. We had observed this phenomenon in our own imaging of mouse brains, but found little more than anecdotal mentions of it in the literature ^8,9,10,11^. To confirm that these dark objects indeed represent cell nuclei with lower autofluorescence intensity than the surrounding lipid-rich brain tissue, we performed a standard 4% PFA perfusion-fixation followed by cryosectioning and nucleus (DAPI) counterstaining. We found the same low-autofluorescent objects, which had an overlap of nearly 100% with nuclear staining (DAPI), confirming that dark objects in STPT indeed represent cell nuclei (Supplemental Fig. S1).

### Population-wide, regionally-resolved exploration of neuroanatomical features

To automatically collect cytoarchitecture data for each brain, we trained a DNN model to detect and segment the nuclei (low-autofluorescent objects) in all brain regions, including those of the highest density, such as the dentate gyrus (DG). Because of computing constraints, we applied the model systematically to segment a subset of the AMBCP dataset comprising 537 brains (Fig. 1A-C and Methods). The model performed with an estimated 97% cell detection accuracy on a test set, with a false positive rate of <0.01 (see Methods) whenever image quality was sufficient (for exclusion criteria of whole brains or certain regions within sections, see Methods). Using detected cells in each section, we obtained a local estimate of the volumetric cell density (see Methods), which, combined with the pixel-wise registration of brain regions provided by the AMBA, allowed us to estimate the average cell density per region for each brain. Similarly, we evaluated the per-region volume of each brain by linear interpolation across all sections (see Methods). In sum, we simultaneously estimated the 3D cell density (*D*) and volume (*V*) of each region for each brain (see Methods). In total, we estimated per-region *D* and *V* for 532 basic regions annotated in the AMBA, which corresponds to levels 6-8 of the region hierarchy.

**Figure 1:**
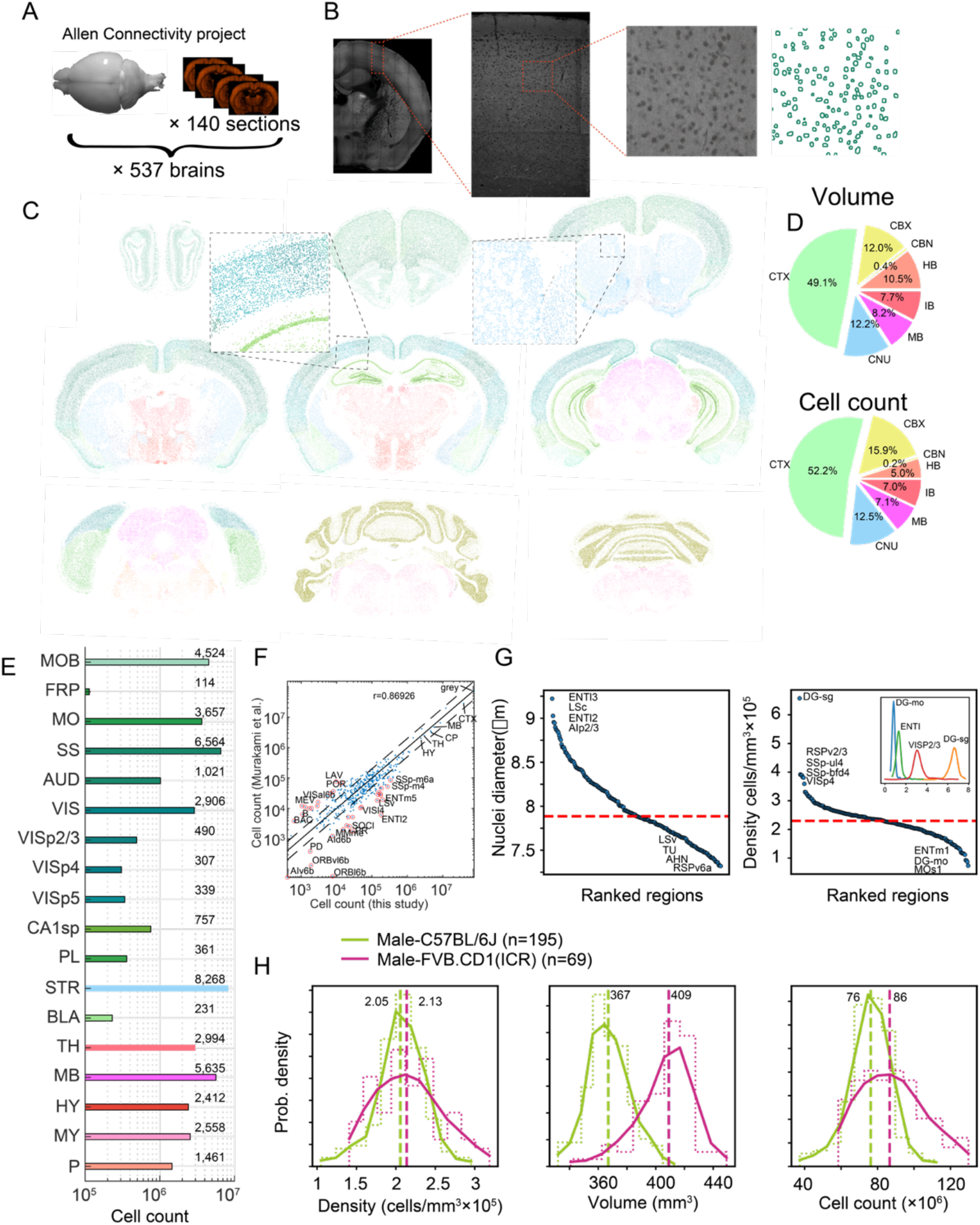
Survey of neuroanatomic properties of the mouse brain. (**A)** The analysis is based on a cohort of 537 mouse brains imaged by serial two-photon tomography using the Allen Mouse Brain Connectivity Project (AMBCP). Each brain comprises ∼140 coronal sections spaced 100μm apart along the anterior-posterior axis. (**B)** Example of nucleus segmentation in the isocortex. Each section was divided into tiles of 312×312 pixels (109×109 um) (zoom-ins, right). A trained deep neural network cell segmentation model (see Methods) was applied to detect the contours of nuclei for downstream analysis across tiles, sections, and whole brains, as shown. **(C)** Segmentation of several sections of one particular brain; segmented nuclei are colored using the Allen Mouse Brain Atlas (AMBA) region convention. **(D)** Pie charts of the median volumes and cell counts across all 537 brains in the main brain regions, colored using AMBA nomenclature. (**E**) Median cell counts for selected brain regions in C57BL/6J males (number near bars in thousands). (**F**) Comparison of region cell counts between this study and Murakami et al., over C57BL/6J males; dots above/below the dashed lines represent regions with greater than two-fold difference. **(G)** Ranking of 532 regions by nucleus diameter (left) and density (right). Each dot corresponds to the median value of one region over 537 brains. Red dashed line, median across regions. Inset shows distributions of density over 537 brains for selected regions. **(H)** Distribution of cell density (left), brain volume (middle), and cell count (right), comparing C57BL/6J males and FVB.CD1 males across basic cell groups and regions (“grey”).

Cell count (*N*) is the product *V* × *D*, therefore it was not considered an independent variable. The median male C57BL/6J mouse brain contained a total of 76 × 10^6^ cells, in 367 mm^3^ of grey matter, at a density of 2.05 × 10^5^ cells/mm^3^. A pie chart of the volume and cell count of the main regions (level 4 of region hierarchy) calculated across 537 brains appear in Fig. 1D, and absolute cell counts for C57BL/6J male mouse representative regions are shown in Fig. 1E. We quantified each level of the hierarchical tree structure of the AMBA and found good correlation (r=0.89) with a recent 3D whole-brain single-cell resolved light-sheet microscopy study^12^ (Fig. 1F). The diameter of detected objects (nuclei) varied between 7-9.5μm (Fig. 1G left), which at a nucleus/soma volumetric ratio of 0.08 ^13,14^ corresponds to median cell body diameters from 16.25μm in the RSPv6a, to 22μm in the ENTl3. The regional variability of cell densities was high, ranging from 1 × 10^5^ mm^-3^ in layer 1 isocortex (e.g., MOs1) to 6 × 10^5^ mm^-3^ in the dentate gyrus granule layer (DG-sg). We show examples of regional distributions across the full cohort of 537 brains in the inset of Fig. 1G right.

The large number of AMBCP brains in our analysis enabled us to compare variabilities of macroscopic properties between subsets of the cohort, e.g., to compare strains. We compared distributions of volume, cell density, and cell count at the coarsest hierarchical atlas level, i.e., across grey matter cell groups in the brains of male C57BL/6J vs. male FVB.CD1 mice (Fig. 1H). Median cell density was similar for the two strains, with considerably larger variance in FVB.CD1 males. FVB.CD1, however, had 11% larger grey matter volume than C57BL/6J. Combining these two features revealed a ∼10% increase in the median cell count in FVB.CD1 vs. C57BL/6J (Fig. 1H right panel). These results suggest that: (*a*) there is no simple relationship between volume and density, therefore, both properties should be measured simultaneously, and (*b*) a large cohort enables detection of relatively small differences.

To test the power of our model, we explored the densities and nucleus diameter of cortical regions (Supplemental Fig. S2). First, we considered the hippocampal formation (HPF) because imaging-based quantification of its denser regions (pyramidal layers of Ammon’s horn and the granule layer of the dentate gyrus) has been difficult^3^ and was achieved only recently ^12,15^. Analyzing 195 C56BL/6J male brains, we found that the pyramidal layer of CA1 was denser than that of CA3 and CA2, whereas nucleus size was larger in CA3. In the dentate gyrus, the granule layer had the highest density of all regions, with >6.5×10^5^ cells/mm^3^, and nuclei were largest in the polymorph layer (Supplemental Fig. S2 upper panels). In the isocortex, we examined the extent to which the cortical layers across cortical divisions differed in density and size (Supplemental Fig. S2 lower panels). Layer 1 was consistently underpopulated, having a density of about 10^5^ cells/mm^3^. The overall rank order from densest to sparsest was maintained, with layer4 > layer6a > layer2/3, layer5 > layer1, suggesting a similarity in cytoarchitecture between cortical regions.

Layer 4 of the primary visual and somatosensory cortices had higher density than did the auditory and visceral cortices. Nucleus diameters showed less distinct distributions between layers, although layer 2/3 and layer 5 tended to have larger nuclei than did layers 4 and 6a.

### Density differences between left and right hemispheres

Brain laterality has been discussed in the literature since Broca and Wernike found language dominance on the left side of the brain. Although evidence for brain laterality in mice is more scarce, a recent study has suggested functional and circuit differences in the auditory cortex^16^. We performed a systematic comparison of the left and right hemispheres seeking differences in region-wise volume and/or cell density. In C57BL/6 males (n=152), we found 67 regions with lateral cell density bias of over 5%, and up to 30%. Both sides showed similar numbers of biased regions in density (L, 37; R, 30, Fig. 2A). Cortical areas VISC2/3 (Fig2b), AUDv2/3 (Fig2c), GU2/3, RSPv6a, and LSc showed more than 20% higher density in the left hemisphere; APr, HATA, ACAd2/3, and PAR (Fig. 2D) were up to 20% denser in the right hemisphere. Strikingly, we found consistent density bias in left hemisphere cortical regions, specifically in layers 2/3. This bias was both most consistent (Fig. 2E) and pronounced (Fig. 2F) in a group of neighboring ventro-lateral areas: visceral, gustatory, temporal association and auditory, where the latter may be consistent with the Levy et al. findings. Laterally biased regions were consistent across strains and sexes (e.g., in females C57BL/6, n=140, Supplemental Fig. S3).

**Figure 2:**
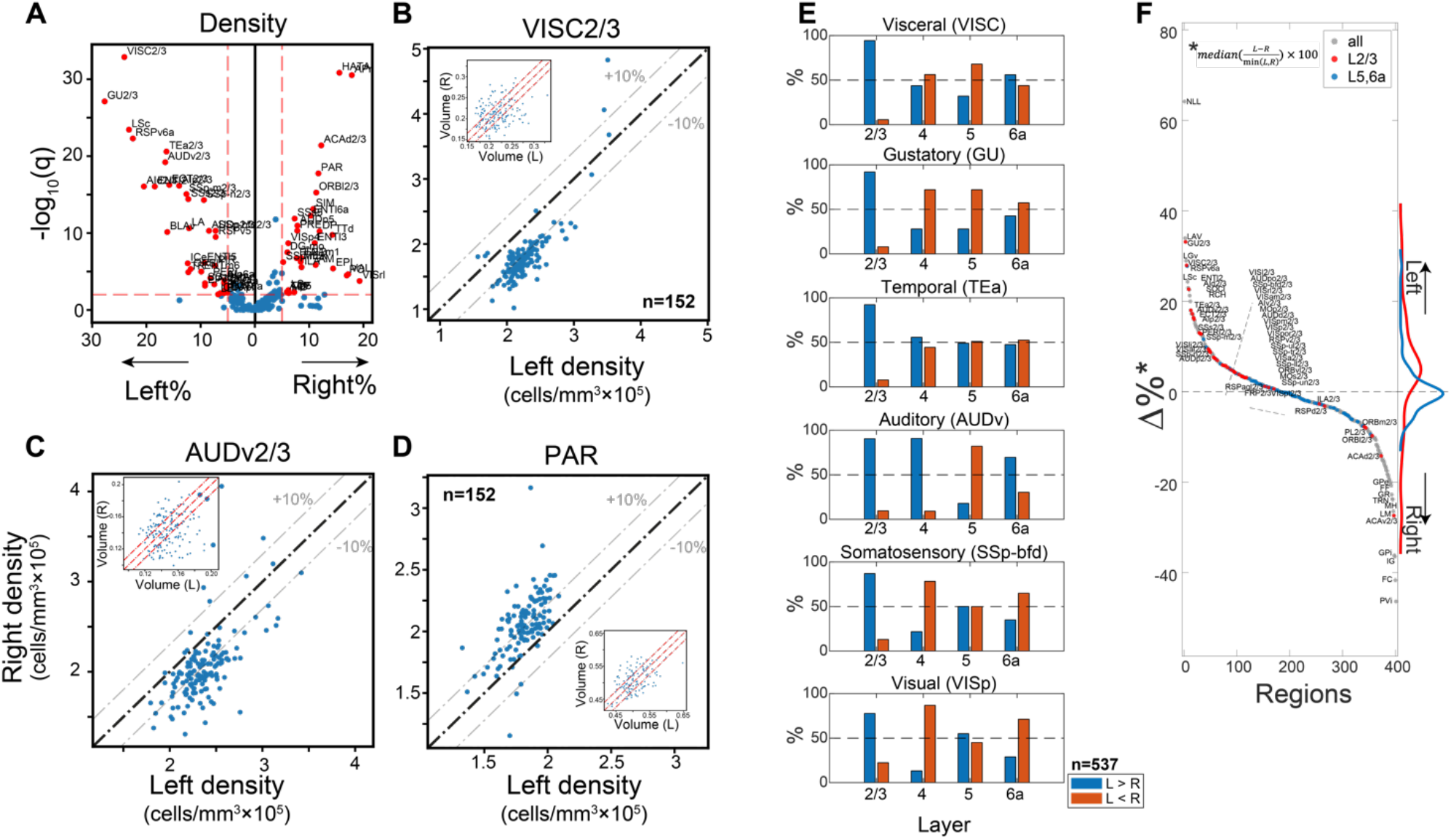
Region-wise laterality of cell densities in C57Bl/6 males. **(A)** Volcano plot comparing average region cell density in left and right hemispheres (x-axis), with rank sum test q-values (y-axis). **(B-D)** Scatter plots of the density of the left vs. right hemisphere for ventral auditory area layer 2/3 **(B)**, visceral area layer 2/3 **(C)**, and parasubiculum **(D)**. Each dot corresponds to one brain (n=152). Insets show corresponding volumes in the same fashion, with no observed lateral bias. **(E)** Percentage of brains with tendency for left or right hemisphere density (blue, left>right; orange, right>left). (**F)** Laterality differences in density (all brains n=537) shown for all regions whose volume >0.05mm^3^ (excluding layer1 and 6b). Regions are sorted by their bias to the left. Red and blue dots show layer 2/3 and layer 5/6a, respectively. Right inset shows the distribution of layer 2/3 regions and layer 5/6a in red and blue lines, respectively.

We note that volume laterality was less pronounced (Supplemental Fig. S3), with some regions slightly larger in the right hemisphere (e.g., PL6a, SSp-ul6a and DG-po; R-L difference ∼8%). Most regions did not show laterality in both density and volume (e.g., Fig. 2B-D display density laterality, while no bias in volume is observed).

### Regions with volume/density sexual dimorphism in C57BL/6J mice

To examine whether differences in overall brain volume or density (Fig. 1H) are isotropic, we conducted a region-specific analysis of volume, density, and cell count. Differences between males and females in regional neuroanatomy have been extensively described, including dimorphic volume and cell count in the medial amygdala (MEA) ^17,18^ and in the bed nuclei of the stria terminalis (BST) ^19^. We first compared C57BL/6J males (n=140) with females (n=152). At the global level (“grey”), males and females had similar total numbers of cells (77 × 10^6^ and 75 × 10^6^, respectively). These similar counts were achieved differently, however: females had a larger median grey matter volume, whereas males had higher median grey matter density (Fig. 3A). We conducted rank sum testing on each region that passed QC (see Methods) for sex differences, in volume and density (Fig. 3B). With the notable exception of both MEA and BST, most regions were consistent with the overall trend of larger volumes in females; many were 5-10% larger. Volume sex differences were compensated by higher cell density in the male brains, leading to slightly more cells in most brain regions in males (see also Supplemental Fig. S4 which shows similar volcano plots for FVB.CD1 mice). We further demonstrated this discordance between median sexual difference in volume vs. density in Fig. 3C, where most brain regions fell in quadrant IV of the volume-density plane. Notable exceptions included the MEA and BST, which were consistently larger in males, and the orbital area layer 2/3, consistently larger in females. Next, we looked beyond the rank sum statistical test, governed by the median of the distribution, at examples of how distributions differ. For example, the ventrolateral orbital area layer 2/3 (ORBvl2/3) showed both larger volumes and slightly higher density in females (Fig. 3D left), resulting in significantly more cells in females (Supplemental Fig. S4A). The opposite was the case for BST, where males had both larger volume and higher density (Fig. 3D middle). As a third example, we showed the case of primary auditory area layer 5 (AUDp5), which displayed no difference in region volume, yet density in the male brains was higher (Fig. 3D right).

**Figure 3:**
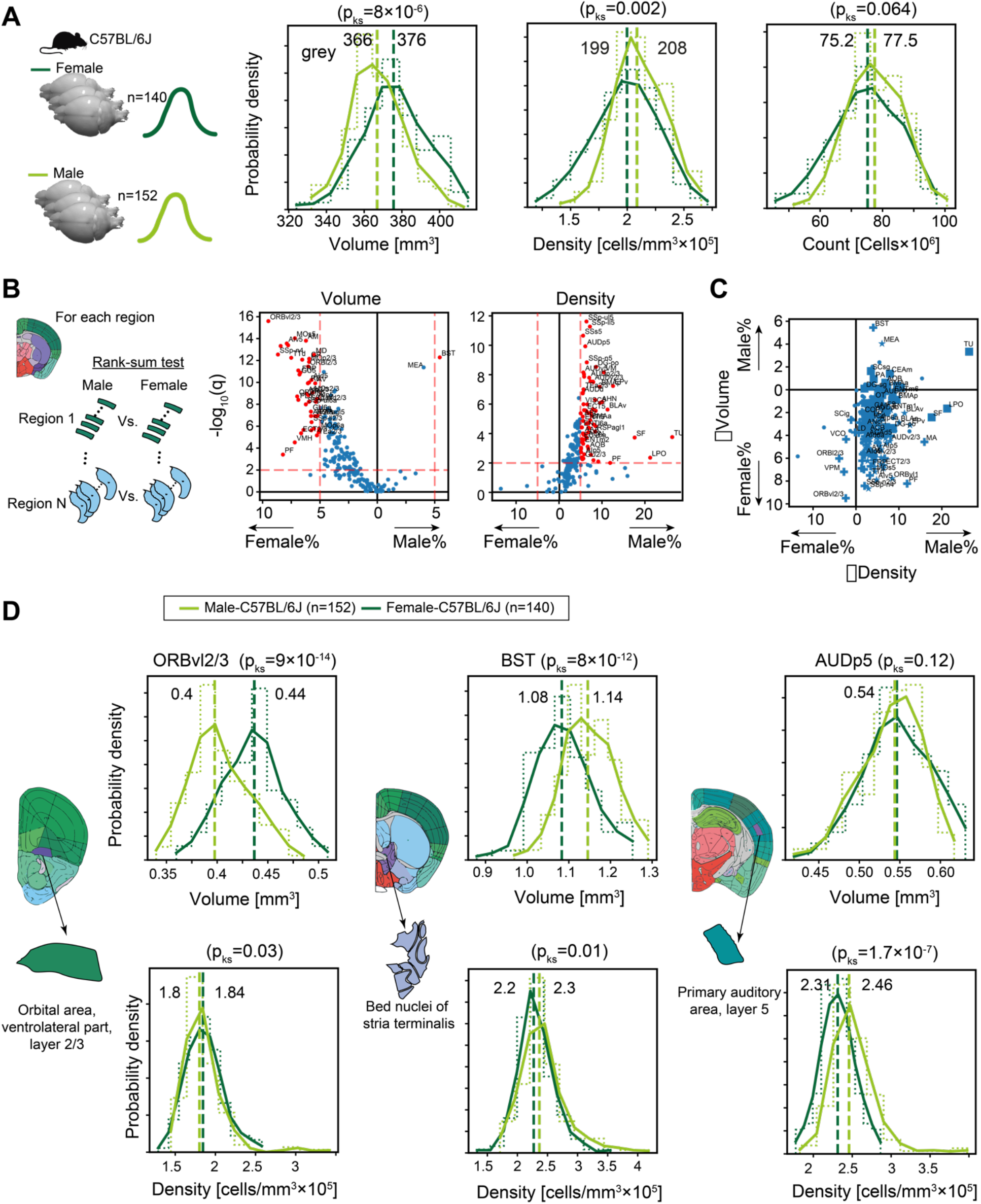
Sexual dimorphism in C57BL/6J. **(A)** Distribution of volume (left), density (middle), and cell count (right) for the whole brain grey matter (“grey”) in female (dark green) and male (light green). P-values correspond to a Kolmogorov-Smirnov test. **(B)** Volcano plots showing per-region statistical testing for male versus female difference in volume (left) and density (right), each dot representing one region. Horizontal axis, median differences (%); vertical axis, q-values (FDR corrected rank-sum p-values by BH procedure in -log10 scale). Red dots highlight regions with an effect size larger than 5% and q<0.01. **(C)** Scatter plot of tested regions (dots), showing median differences in volume vs. density. Markers represent statistical significance: both volume and density (star), volume only (+), or density only (square). **(D)** Examples of regions that display sexually dimorphic volume and/or density. Distributions of volumes appear in the upper row, distributions of densities in the lower row.

In sum, the population-wide survey revealed a number of sexually dimorphic areas; Yet, to what extent were region volumes or densities predictors of the individual’s sex? To answer this question, we trained linear support vector machine (SVM) classifiers. In brief, we randomly selected 2/3 of the C57BL/6J brains (n=184), where each brain was represented by a 532-dimensional vector of either volume or density across regions, and trained the SVM. We then tested its performance on the remaining 1/3 of brains (n=97). This process was repeated 100 times and resulted in an average accuracy of 78% and 90% for density and volume, respectively. From this, we identified the regions that had the highest contribution to the separating hyperplane (Fig S4e, g), and trained new classifiers based on these top regions, adding one region at a time. Using volume data, MEA and BST alone performed classification to >70% accuracy. Adding the rostral lateral septal nucleus (LSr), the classifier’s performance saturated to about 90% (Suppl. Fig. S4F). Suppl. Figure S4I shows the SVM separating line based on MEA and LSr yielding an accuracy of 84%. In contrast, the density-based classifier showed only incremental improvements when adding almost any of the top 10 regions (Suppl. Fig. S4H). Together, these results suggest that although many brain regions show significant differences between males and females both in volume and density (Fig. 3), the overlap between the male and female distributions in each of these regions hampers classification. Apart from MEA and BST, the classifier revealed the rostral aspect of the lateral septal nucleus (LSr) as a strong predictor for sex in volume and density (higher in female), in addition to e.g., olfactory (PA, TR, COA) and prefrontal (ILA, ORB) areas.

### Strain differences in volume and density

Following the observation in C57BL/6J mice that female brain volumes were higher despite a smaller body size, we investigated the relation between recorded body weight and grey matter volume. To this end, we added the cohort of outbred FVB.CD1 mice, a strain with 40-50% higher body weight than C57BL/6J. As expected, in both strains, males and females showed distinct distributions for body weight, and males were larger than females (Fig. 4A). Distributions for grey matter volume had higher overlap between sexes and showed opposing trends between the strains: in contrast to C57BL/6J, FVB.CD1 females had smaller brain volumes than males. Moreover, within each strain, body weight did not correlate with grey volume. We next quantified sex and strain differences in brain volume and density, resolved to neuroanatomical regions. First, we compared strain differences in females with those in males, showing concordance/discordance patterns between males and females (sex) (Fig. 4B-C and schematic to the right). Second, we compared sex differences in FVB.CD1 with those in C57BL/6J, showing concordance/discordance patterns between strains (Fig. 4D-E and schematic to the right). *Strain-wise analysis*: FVB.CD1 brains were overall larger, but the volume expansion with respect to C57BL/6J was not uniform across regions. Region volumes ranged up to 30% differences, with the extreme example of the cerebellum (CENT2), whose size increased by 50% in both FVB.CD1 males and females (Fig. 4B). Moreover, per-region volume differences between strains were, in general, larger in males (i.e., most data points in Fig. 4B quadrant III are above the diagonal). Only two regions showed larger volumes in C57BL/6J: the main olfactory bulb (MOB) and the caudal lateral septal nucleus (LSc) (Fig. 4B quadrant I).

**Figure 4:**
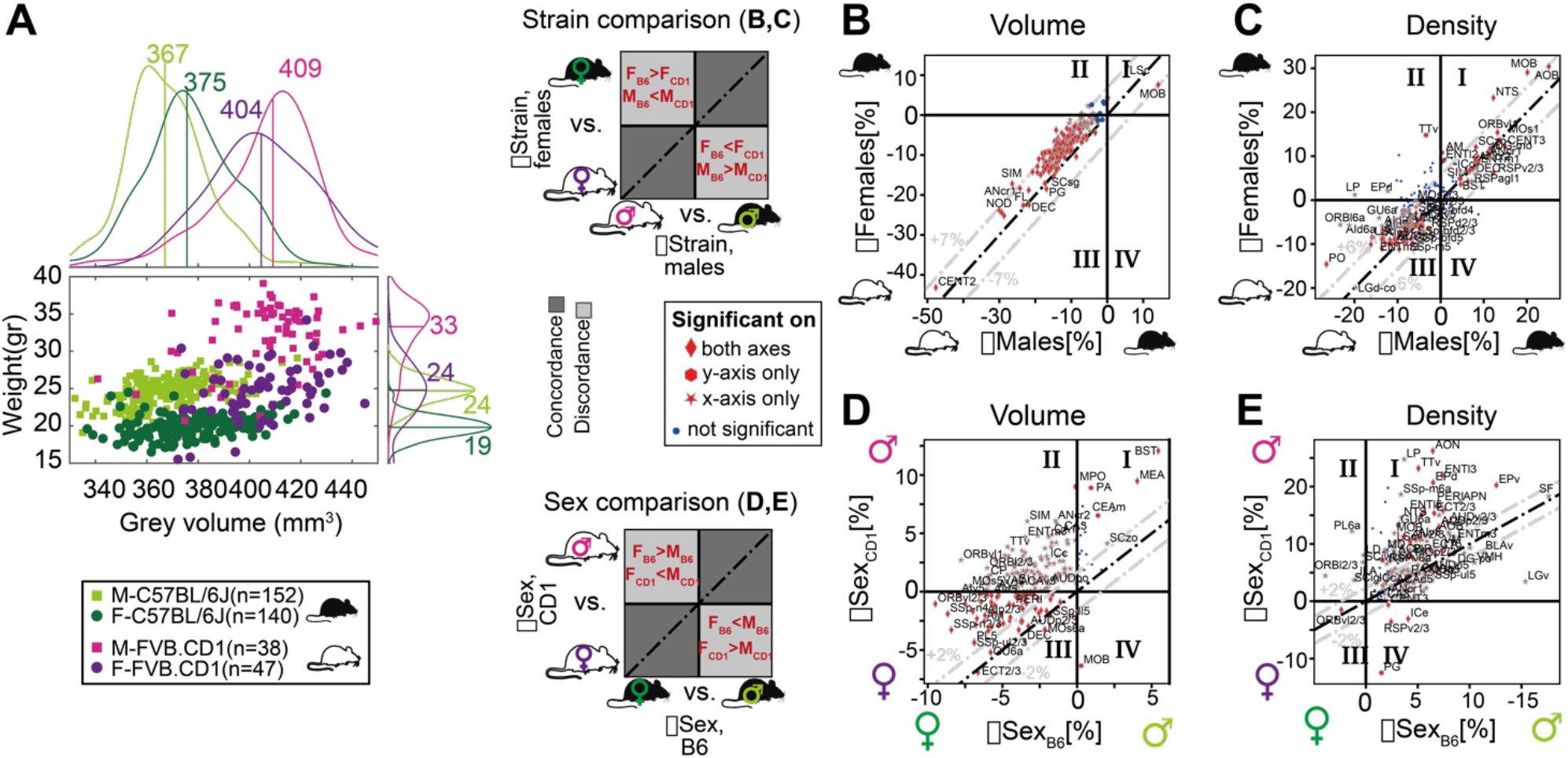
Sexual and cross-strain dimorphism in C57BL/6J (B6) and FVB.CD1 (CD1). **(A)** Scatter plot showing body weight vs. grey volume for 537 brains. Side panels show the group distributions of grey matter volume (upper) and weight (right). Lines are the medians whose values are indicated. **(B-C)** Strain comparison of per region volume **(B)** and density **(C)**. Differences between the median values of the strains, per region, are shown for males (horizontal axis) and females (vertical axis). Points in quadrants I and III suggest concordance between males and females across strains, as illustrated in the schematic on the left. Red markers designate statistical significance in either axes or in both. (**D-E**) Sex comparison of per region volume **(D)** and density **I**. Points in quadrants I and III suggest concordance between C57BL/6J and FVB.CD1 across sex, as illustrated in the schematic on the left.

A similar comparison for cell density per region suggests non-uniform density differences, with almost half the regions being denser in C57BL/6J, and the other half in FVB.CD1 (Fig. 4B quadrants I and III, respectively). In this comparison, olfaction-related regions (AOB and MOB) showed higher density in C57BL/6J, while the LSc showed the opposite effect.

*Sex-wise analysis*: Differences in volume confirmed sexual dimorphism in MEA and BST, which were larger in males for both strains. These differences were more pronounced in FVB.CD1 than in C57BL/6J (Fig. 4D quadrant I). Many brain regions showed “strain-discordant” dimorphism, with females having a larger volume in C57BL/6J and males in FVB.CD1 (Fig. 4D quadrant II). Although total brain volume in FVB.CD1 males was larger, some regions showed larger volume in females (e.g., the previously mentioned orbital cortex ORB, Fig. 4D quadrant III). Comparing sexual dimorphism in density (Fig. 4E), we found a simpler and more consistent picture: in both strains, males had higher density in all regions except for ORBvl2/3. Note that in density as well, sex differences were found to be larger in FVB.CD1 (most data points in Fig. 4E quadrant I are above the diagonal).

### Region-wise correlations between volume and density across brains

To the best of our knowledge, no previous study simultaneously quantified cell density (*D*) and brain region volume (*V*). We therefore sought to investigate whether constraints exist between *D* and *V*. For example, if the number of cells in a region is constant across brains, *D* and *V* must be negatively correlated. If, by contrast, the number of cells in a region, *N*, scales with the volume while *D* remains constant, *D* and *V* display zero correlation. If a positive correlation exists between *D* and *V*, the number of cells *N* grows faster than linear with respect to either *D* or *V* (Fig. 5A). Based on per-region measurements of both *V* and *D*, we calculated regionally-resolved Pearson correlations between volume and density (Fig. 5B). In 72% of regions (289/397), cell density was negatively correlated with volume (Fig. 5C), with a median correlation of −0.096. For example, we showed two regions where *N* was positively correlated with both *D* and *V*, yet the correlation between *D* and *V* was either positive (AAA, Fig. 5D) or negative (SSs2/3, Fig. 5E). This suggests that for some regions, cell count does not scale simply or linearly with volume.

**Figure 5:**
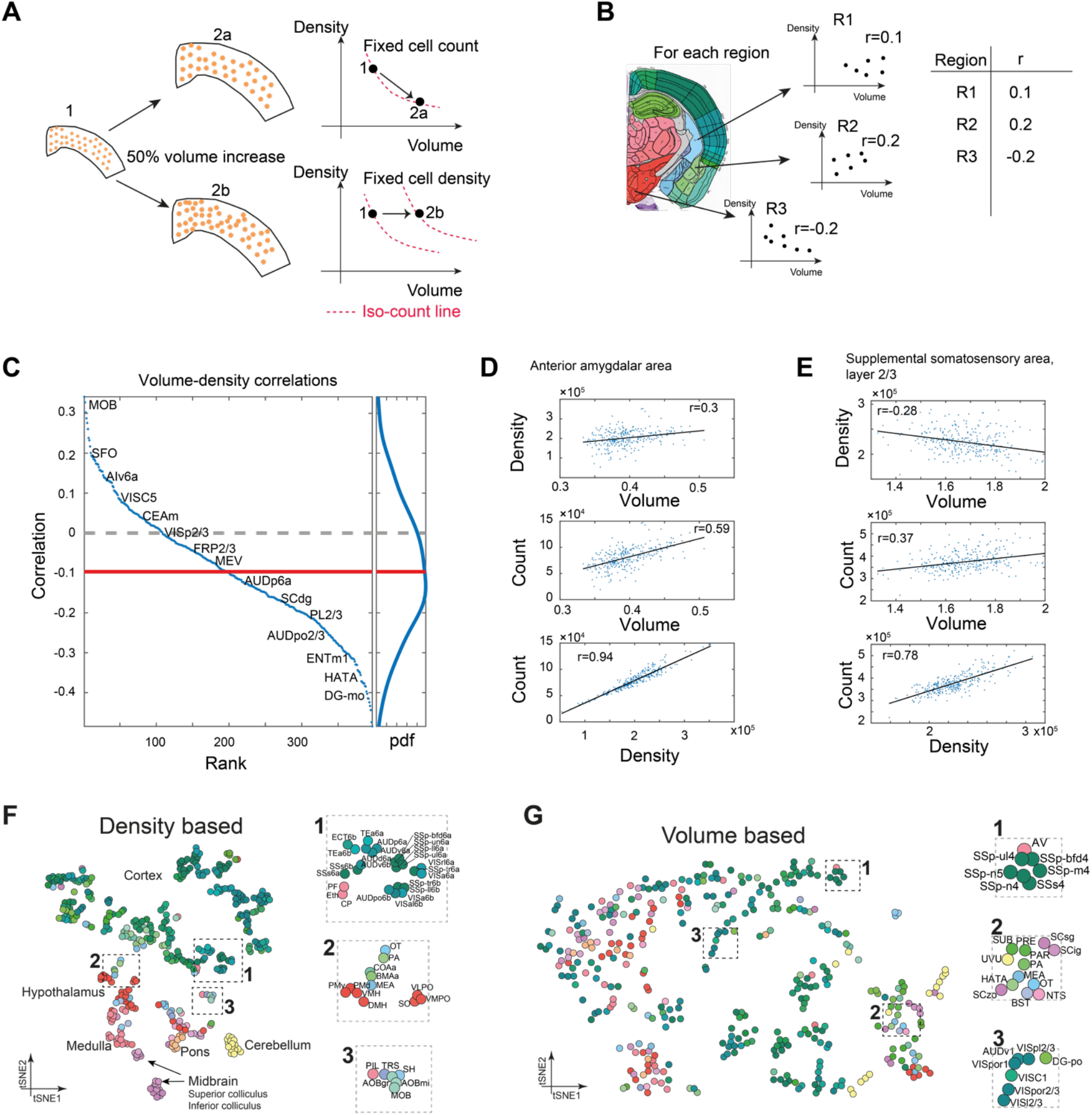
Correlations between volume, cell count, and density. **(A)** Schematic illustration of two types of relations between regional cell density and volume, associating region expansion with a fixed number of cells (upper) or with a fixed density (lower). Each regional expansion can be represented by a shift in the volume-density plane (right column). **(B)** A scheme showing how for each region the correlation between density and volume was measured over the whole dataset. **(C)** Brain regions ranked by the correlation between volume and density. Correlations higher than 0.13 or lower than −0.13 correspond to q-values lower than 0.05. Side panel displays the distribution of correlation values, and its median is denoted by the red line. **(D)** Correlations between volume, density, and count in the anterior amygdalar area. **(E)** As **(D)**, for supplemental somatosensory area, layer 2/3. **(F-G)** Visualizing similarity between brain regions across the population, by tSNE embedding on pairwise correlations between region density (**F)** or region volume **(G)**. Each dot represents a region and is colored according to the AMBA convention. Dashed rectangles indicate zoom-ins on three frames on each tSNE.

### Inter-brain similarity between regions based on volume and density

Finally, we assessed similarity between regions, based on volume or density. We used tSNE as a 2D embedding method over the density data (Fig. 5F-G). Briefly, each region is characterized by a vector of 537 components, each representing its density across one brain. 2D embedding aims to preserve the local similarity between regions.

The density-based tSNE embedding map in Fig. 5F reveals clear 2D “clusters,” largely consistent with neuroanatomical classification. Cortical regions appear in the upper part of the map (colored green), and cerebellum (yellow), midbrain, and hindbrain in the lower part. We further explored whether the order within the cortical part may be explained by layer structure or by cortical division, but found no clear structure (Supplemental Fig. S5a-b). Compared to Fig. 5F, tSNE embedding based on volume was more “dispersed” and displayed disorder with respect to neuroanatomical classification (Fig. 5G). To demonstrate that the tSNE map is indeed based on true variations in region-to-region correlations, we compared density-based and volume-based correlations. First, for each region we identified its 10 most correlated regions based on either density or volume. These correlation values were higher for density-based correlations across almost all regions (Supplemental Fig. S5C), supporting the observed density-based “order.” Second, we selected 10 representative regions across the brain and calculated all their pairwise correlations (Supplemental Fig. S5D), showing that even for distant regions, density-based correlations remain much higher than volume-based correlations. Thus, similarity in volume across regions is less “preserved” brain-wide than similarity in cell density.

## Discussion

We presented an automated, imaging-based, staining-free study of neuroanatomy and cytoarchitecture in the mouse brain. We conducted our measurements on a massive, high-quality dataset of serial two-photon tomography ^6^, aligned with a well-annotated reference atlas ^8^.This made possible, for the first time, a detailed population-wide analysis of two important neuroanatomical variables simultaneously: cell density and volume, resolved for 532 regions. The data spans an unprecedented cohort of 537 mice of two strains, the inbred C57BL/6J, and the hybrid FVB1.CD-1, each represented by both females and males. Our high-throughput measurements of cell densities were achieved by using a DNN trained to detect low-autofluorescent cell nuclei with high accuracy, even in the most cell-dense regions of the brain.

The study has several limitations. First, the model is sensitive to image quality, and in particular, contrast between dark nuclei and autofluorescent surroundings. In the hindbrain (pons, medulla), contrast was exceedingly weak, and we expect our quantifications in this region to strongly underestimate real cell densities, to an extent we cannot quantify. Second, AMBA annotations were not always resolved to the most refined level of the atlas hierarchy. For example, density values for the cerebellum appear to be uncharacteristic because the cell-dense granule layer and sparse molecular layer were not distinguished at the deepest level of annotation (e.g., CENT3 included the granular and molecular layers). The same is true for the hippocampus CA1-2-3, where we used cell density-based clustering to distinguish the pyramidal layer (sp) from its surrounding sparse layers (slm, so, sr, see Methods). Therefore, although the model performed exceedingly well even in these cell-dense regions, the absence of annotations stood occasionally in the way of making biologically meaningful distinctions.

Nevertheless, we provided key statistics that help answer fundamental, recurring questions in neuroanatomy. Although no other study presented simultaneous measurements of volume and cell density, our data correlate well with a wealth of literature in the field. We achieved good region-wise correlation with full 3D volumetric cell counts by expansion microscopy ^12^ (Fig 1F). Our derived cell count of mouse brain grey matter (76 × 10^6^ for male C57BL/6J) is well within the range of existing cell count estimates for adult males (6 × 10^6^cells) ^20,21,12,15^.

By measuring the largest cohort to date, we provided partial support for the notion that this extreme range in the literature may not stem from variation in strain or sex, but rather from individual differences ^12^. The median cell counts between sexes and strains differed no more than 13% overall, or ∼40% for the most deviant individual structures (MOB and CENT2), while the standard deviation across individuals was ∼10 million cells (for C57BL/6J male alone). Hence, values between 55 × 10^6^ − 95 × 10^6^ are within ±2 standard deviations of the distribution of total grey matter cell counts. We claim that the notion of “ground truth” values of brain cell number *can* be reached, yet are best reflected by a population distribution.

Our dataset provides a large, important corpus toward this “ground truth,” and similar studies can further help distinguish technical biases from biological variation. For example, our cohort provides the first brain-wide survey of cell architecture laterality, with numerous regions showing density bias to the right or left hemisphere; and some volume bias in favor of the right hemisphere. We describe a remarkable phenomenon of ventro-lateral cortical areas increasing cell density in left hemisphere layer 2/3, but no other layers, that may be interesting to follow up functionally.

We further validated, and expanded on the existing literature describing examples of regions that show sex- or strain-based differences. Sexual dimorphism of brain regions was significant, as exemplified by the success of our SVM-classifier, yet the ability to separate males from females was limited due to population’s interindividual differences. For instance, medial amygdala and bed nuclei of the stria terminalis were both larger and denser in males, but to a lesser extent than reported in smaller studies ^22^ and to a similar extent to what was reported in MRI-based studies ^23^. By contrast, in females, several prefrontal cortex structures were larger (e.g., ORBvl2/3), which resulted in higher cell counts. We found no evidence of this phenomenon in the literature on mice, but an MRI population study of 2,838 human individuals found higher grey matter volume (GMV) in prefrontal areas in women ^24^. Between the strains, we found considerable differences in the olfactory system, which was larger and denser in C57BL/6J, and in the cerebellum, which was larger in FVB.CD-1. Finally, we provide an accessible, web-based platform for open exploration of the data. The web application allows researchers to freely compute distributions of any measured neuroanatomical features, for any brain region, and across the entire population or specific subsets. This exploratory resource can be of great use for experimental design, and lead to more accurate brain modeling.

## Online Content

### Code Availability

Segmentation-related code is available at https://github.com/delkind/detectron_segmentation.

### Brain Explorer: data viewer for mouse brain cell data

We provide a tool for examining analyzed AMBCP brain images. The tool allows visualizing distributions of macroscopic parameters (i.e., density, volume, cell count etc.) for any region across any subset of the brains, as well as performing statistical tests. Subsets of brains can be based on any combination of strain and sex. Resulting charts can be exported as CSV or PDF files. The viewer is available in https://github.com/delkind/mouse-brain-cell-counting.

## Acknowledgements

N.S. is funded by the Ministry of Science, Technology & Space, Israel (Grant 3-16033). A.Z. is supported by European Research Council (TYPEWIRE-852786), Human Frontiers Science Program (CDA-0039/2019-C) and Israel Science Foundation (2028912). H.H. is supported by the Swedish Brain Foundation (Hjärnfonden) and Human Frontiers Science Program (CDA-0039/2019-C). We thank Eytan Domany for critical discussion. We thank Dvir Aran and the Open University for computing resources.

## Author contributions

A.Z., N.S. and H.H. conceptualized and designed the study. D.E., N.S. and A.Z. curated and analyzed all data. D.E. developed and implemented all code and tested all software with N.S. and A.Z. D.E., H.H., N.S. and A.Z. critically discussed and interpreted data. E.A. collected brains and performed histological validations. H.H., A.Z. and N.S. designed figures and wrote the paper with help from D.E. and E.A.

## Methods

### Data

The Allen Mouse Brain Connectivity Project (AMBCP) dataset ^6^ consists of 2,992 brains, of which we processed 537 and eventually used 399 in our analysis (the strain and sex breakdown of the brains appear in Table 1). Each brain consisted of ∼140 section images captured every 100*μm* along the anterior-posterior axis using two-photon tomography ^5^. Image resolution was 0.35*μm* per pixel. AMBCP post-processed section images for noise removal. Rather than using the red, green, and blue channels that display brain connectivity, we used the background channel of the images, as provided by AMBCP, without additional processing, except for converting the RGB image to grayscale.

**Table 1:**
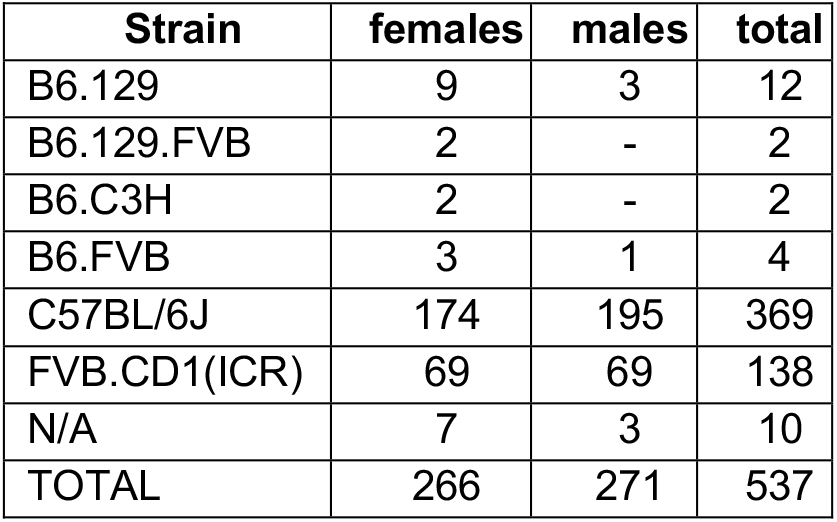
Breakdown of the data by strain and sex

| Strain | females | males | total |
| --- | --- | --- | --- |
| B6.129 | 9 | 3 | 12 |
| B6.129.FVB | 2 | - | 2 |
| B6.C3H | 2 | - | 2 |
| B6.FVB | 3 | 1 | 4 |
| C57BL/6J | 174 | 195 | 369 |
| FVB.CD1(ICR) | 69 | 69 | 138 |
| N/A | 7 | 3 | 10 |
| TOTAL | 266 | 271 | 537 |

### Training a deep neural network for cell segmentation

To detect cells in an image and mark their contour, we used the Detectron2 deep neural network library ^25^, which relies on a Mask R-CNN image segmentation model ^26^ with the ResNet-101 ^27^ as its backbone.

### Model training and validation

Training the model required 3 rounds of manual annotation and training.

#### Initial manual annotation of the data set and model training

We annotated cell contours manually using the VGG Image Annotator software ^28^. Initially, we annotated only the hippocampus, which is relatively large and easily discernible. The hippocampus contains sub-regions of different densities, which we believed would adequately represent the variety of cell densities across the mouse brain. We manually annotated tiles of 312 × 312 pixels (109 × 109*μm*), randomly selected from the hippocampus in 5 sections of 3 brains (55 tiles in total). We provided these tiles to the network as training data, together with basic data augmentation (e.g., rotation and brightness changes) ^29^.

#### Retraining on hippocampal sections

We then applied the trained model to detect cells on a new set of 55 randomly selected hippocampus tiles. We manually corrected the results produced by the network to create a new set of ground truth annotations. Next, we retrained the model from scratch over a combined training set of 110 tiles.

#### Retraining on other regions

We subsequently used the trained model to detect cells on random sections of 3 selected brains. Visual inspection enabled us to select a set of 64 tiles that displayed the least accurate results and annotate them manually.

#### Final training

We retrained the model from scratch on the resulting training set of 174 tiles (selected from ∼15 sections of ∼10 brains). The total number of cells across the training set tiles was 6,247, corresponding to 0.008% of the estimated 77 million cells in the whole brain.

#### Technical details

We conducted the training with a batch of size 2, a learning rate of 0.00025, with decay, using the Adam optimizer ^30^. Training over 174 tiles required ∼395,000 iterations, and took ∼36 hours using a Linux server with 160 Intel Xeon Gold 6248 2.5GHz CPUs and a Tesla V100S-PCIE-32GB GPU.

#### Evaluating model performance

The training process completed when the model converged. The accuracy of the model on the training data was 99.8%, with a false negative rate of 0.4%. To evaluate model performance, we manually annotated 30 additional tiles from the isocortex, medial amygdala (MEA), hypothalamus (HY), and hippocampus (HIP) of 27 brains and compared them with model prediction (Table 2). We obtained highly accurate results, comparable to the performance over the training data, for segmentation scores such as Jaccard measure ^31^, F1 score (harmonic mean of precision and recall), and total errors (i.e., percentage of mislabeled pixels), as well as for detection scores such as accuracy (detected cells divided by total cells) and false positive rate (false positives divided by total cells).

**Table 2:** Model performance over out-of-sample tiles

|  |  | Segmentation (pixelwise) scores |  |  | Detection (cellwise) scores |  |
| --- | --- | --- | --- | --- | --- | --- |
| Region | # cells in the test set | Jaccard index | F1 | Total errors | Accuracy | False positive rate |
| Isocortex | 192 | 0.982 | 0.991 | 0.002 | 0.962 | 0 |
| MEA | 115 | 0.975 | 0.987 | 0.001 | 0.962 | 0 |
| HY | 163 | 0.953 | 0.974 | 0.003 | 0.938 | 0.005 |
| HIP | 566 | 0.986 | 0.992 | 0.001 | 0.979 | 0.009 |

### Brain-wide automatic segmentation

The trained DNN was applied to 537 brains, as described in detail below.

#### Extracting cell information per section

We divided each section into overlapping tiles sized 312 × 312 pixels, with an overlap of 20 pixels on each side (thus mitigating potential artifacts close to the borders of the tiles). We then applied the trained DNN to detect cells in each tile, resulting in a cell mask (i.e., a Boolean 312 × 312 matrix whose entries are *true* if the corresponding pixel is part of a detected cell and *false* otherwise). Next, we stitched the tiles together using a logical OR over overlapping areas, resulting in a single cell mask per section. Subsequently, we performed contour detection to obtain the coordinates of each cell in a section, and computed the morphological properties of each cell (i.e., circumference, diameter, and area). Following this analysis step, each section image was represented by a table containing the coordinates and morphological properties of its cells.

#### Assigning cells to regions

We used the Allen Mouse Brain Atlas (AMBA) ^8^ to assign the coordinates of detected cells in each section to their corresponding brain region (Table S1). But the atlas annotation was too coarse for several regions of interest, i.e., CA1, CA2, and CA3 of the hippocampus. The common denominator of these regions was the presence of a dense and a sparse region that were not separated by the atlas (e.g., the pyramidal and stratum regions of CA1, CA2, and CA3). To provide the coordinates of these sub-regions, we defined a local measure of density referred to as cell “coverage,” and used it to cluster the relevant cells into a dense and a sparse region. Briefly, in a window of 64 × 64 pixels centered around each cell we counted the number of pixels that belong to cells, thus assigning a local “coverage” measure (the median cell area was 80 pixels, much smaller than the window around it). We then detected the sub-regions by clustering the cells according to their “coverage” values. For example, we took the “coverage” values of all CA1 cells and used K-means clustering to split them into two clusters of high and low “coverage” values. In this way, the coordinate of each cell center was assigned to either cluster. We then drew the circumference of the sub-regions by applying a standard morphological closing operation, and discarded spurious small regions.

#### Estimating volumes, 3D densities, and cell counts

Until this stage, the analysis provided local, i.e., microscopic properties for each detected cell, and assigned cells to a brain region. The next step was to collect cells that belong to each region and estimate their density, the volume of the region, and the total cell count. This required calculating 3D estimates based on the relevant 2D data, using the following steps:

##### (1) Estimating cell density per section

We used AMBA to label the area of a given region in a section. We assumed that cells belonging to a region are equi-radius spheres whose projection on the 2D section depends on the distance between their centers and the optical plane, and on the optical depth of field (Fig. M1). Hence, detected cells on a 2D section *s* originate from a slab whose volume is *v*_*s*_ = *a*_*s*_ · (2 *R* + *d*), where *a*_*s*_ is the area of a region, *R* is the radius of the cells in the region, and *d* is the optical depth of field. Cell density per section, *ρ*_*s*_, is given by dividing the number of detected cells by *v*_*s*_. The value of *a*_*s*_ is measured by pixels whose size is 0.35*μm*, and *d* = 1.5*μm* ^5 32^. The value of *R* was taken as the 90^th^ percentile of measured cell radii in *a*_*s*_. The distribution of cell radii corresponds to the “projection” of the cells on the measured section, together with the optical depth of field. Downstream results of cell count and density significantly depend of the value of *R*, e.g., using the 50^th^ percentile would provide larger estimated cell counts. Yet, rank order of cell counts and densities across regions is independent of the selected value of *R*.

##### (2) Calculating region volume

AMBA provides pixel-wise region annotation for each section, making it possible to calculate the area of a region per section (which is independent of cell segmentation). The 3D volume of a region is given by the sum of region volumes between adjacent sections, estimated by the average of its areas over each section. For example, if a region appears in sections 1, 2, 3, and 4, its volume is the sum of average volumes between sections 1 and 2, 2 and 3, and 3 and 4.

##### (3) Calculating cell counts across adjacent sections and in total

Cell counts between the adjacent sections of each region are given by the average densities in those slides multiplied by the volume of the region between these sections. The total cell count of a region is provided by a sum across all relevant sections.

##### (4) Calculating cell densities per region

The overall density of each region is given by the total cell count divided by the volume of the region.

**Figure M1:**
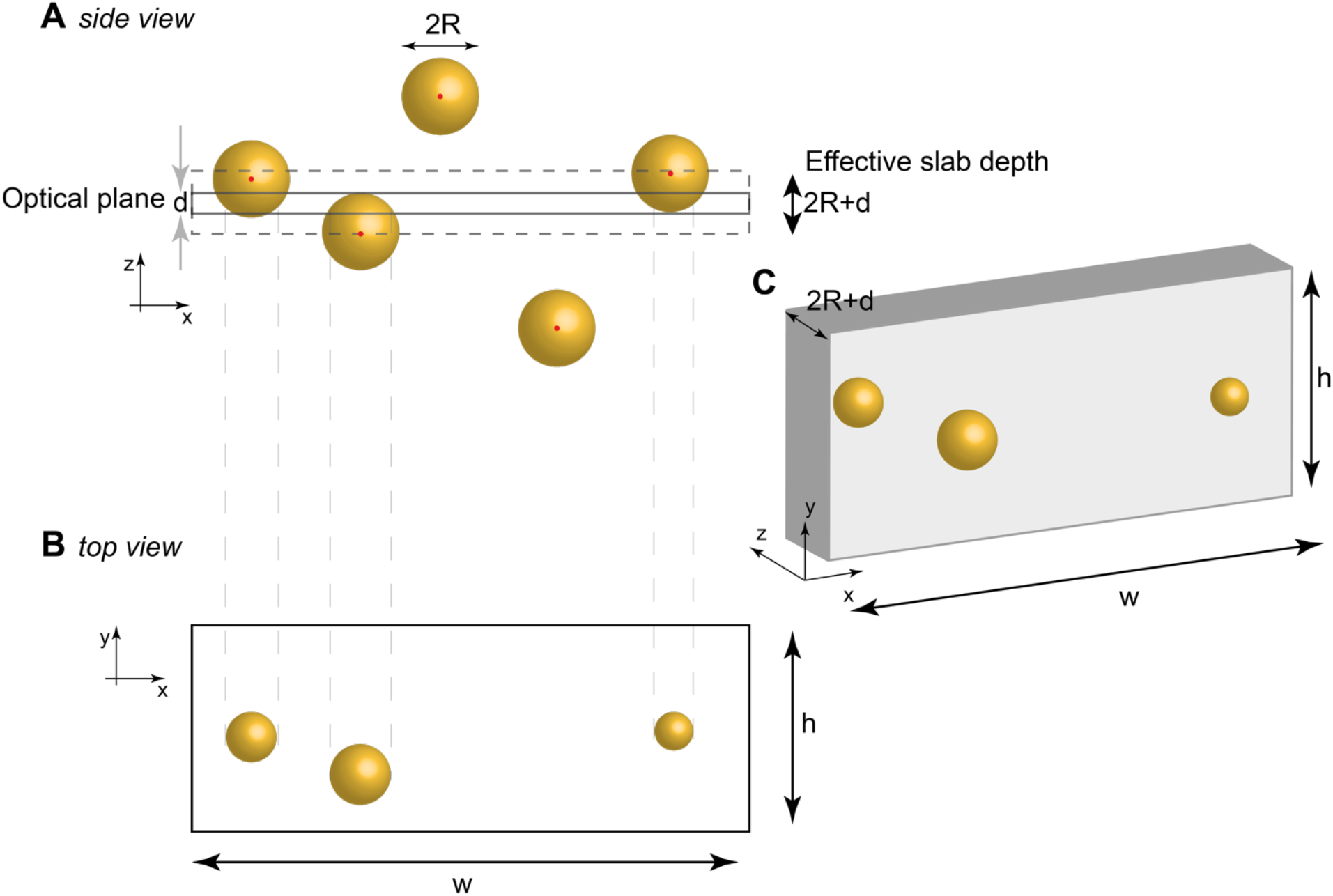
Cell projection onto the section and the relevant volume. (**A**) Cells in 3D vs. the section plane. (**B**) Cell projections onto the plane of the section. Cell whose center is more than ***R*** + ***d*/2** away from the plane are not counted. (**C**) The resulting slab for the purpose of density calculation. The slab volume is ***wh*(2*R*** + ***d*)**, hence the density is **3**/[***wh*(2*R*** + ***d*)**].

#### Discarding whole brains or particular regions of lower technical quality

After calculating the three-dimensional counts and densities across all regions in all brains, we excluded from subsequent analysis regions and whole brains that displayed potentially flawed estimates. We applied the following criteria:

##### We discarded brains displaying dark images

We filtered out brains whose median brightness across the whole brain (“grey” region) was lower than 25 (on a scale between 0 and 255). In such cases, all ∼140 sections of the brain were excluded from downstream analysis because DNN cell detection was either impossible or provided significantly lower estimates.

##### We discarded brains displaying outliers in cell count

We noticed that a common optical artifact of resolution degradation caused the DNN to falsely detect large amounts of excess cells. We marked cases in which cell count in a region was 3 standard deviations larger than the median for the region across brains (calculated as *MAD*. 1.4826, assuming normal distribution). We discarded brains that included more than three such outlier regions.

##### We discarded regions of small volume

We filtered out regions whose median volume across brains was smaller than 0.3*mm*^3^, or whose median cell count across sections was smaller than 500. We excluded such regions across all brains.

##### We discarded regions displaying a correlation between cell count and image brightness

We excluded regions exhibiting strong correlation (>0.25) between brightness and cell count because we assumed that in this case cell count was affected by the inability of the model to discern the cells when the brightness was too low. We discarded such regions from all brains.

##### We discarded regions displaying different estimates in right vs. left hemispheres

Cell count estimates in the right and left hemispheres served as a proxy for technical noise. We calculated cell counts per region using each hemisphere independently. If the difference in cell count between hemispheres for a specific brain was higher than 15.5% of the total cell count for that region, we excluded the case from downstream analysis.

Examples of excluded regions and brains appear in Fig. M1. In sum, we processed 537 brains, of which 138 were fully discarded. Of 690 regions in AMBA, 369 were discarded completely. Across the remaining 399 brains and 321 regions, there were 12,016 (9%) cases in which a region was excluded.

### Predicting sex based on brain region volume or density

#### Performance estimates based on a linear SVM

We randomly selected 2/3 of the C57BL/6J brains (n=184), where each brain is represented by a 532 dimensional vector of either volume or density across regions, and trained the SVM (C=XX). We then tested its performance on the left out 1/3 of brains (n=97). These random splitting into a training and test sets was repeated 100 times. The performance was 0.910±0.022 and 0.780±0.039, for volume and density, respectively.

#### SVM based on specific regions

Each of the regions was ranked according to the average absolute value of its weight in the separating hyperplane across the 100 SVMs, for either volume or density. The top 10 regions (i.e., SVM features) were selected and an SVM was trained on the whole dataset (i.e., without a train-test split) while adding one region at a time.

### Mice

Adult, male C57Bl/6JOlaHsd wildtype mice (Envigo) were housed under standard conditions and provided chow and water *ad libitum*. Experimental procedures followed the legislation under the Israel Ministry of Health - Animal Experiments Council and were approved by the institutional Animal Experiments Ethics Committees at Technion Israel Institute of Technology.

### Histology

Mice were sacrificed by an overdose of Ketamine/Xylazine, followed by transcardial perfusion with PBS, then 4% PFA in 1xPBS. Brains were extracted, post-fixed 24h in 4%PFA at 4°C, cryoprotected in 15%, then 30% sucrose, and finally OCT-embedded, frozen and cryosectioned (20μm). Sections were collected free-floating to PBS. For staining, we first incubated sections 1h in blocking buffer (5% normal goat serum, 3%BSA, 0.3% Triton X-100, in PBS with 0.05% NaAzide), followed by overnight incubation in 1:500 anti-NeuN-PE (Milli-Mark A60, #FCMAB317PE), in the same blocking buffer. After 3 10min washes in PBS, we counterstained 3min with DAPI (NucBlue, Thermo #R37606), mounted and coverslipped sections (Fluoromount-G, invitrogen #00-4958-02), and imaged on a Nikon Ti2 Eclipse inverted fluorescence microscope (20x objective).

## References

1. White, J.G., Southgate, E., Thomson, J.N., and Brenner, S. (1986). The structure of the nervous system of the nematode Caenorhabditis elegans. Philos. Trans. R. Soc. Lond. B. Biol. Sci. 314, 1–340.

2. Keller, D., Erö, C., and Markram, H. (2018). Cell Densities in the Mouse Brain: A Systematic Review. Front. Neuroanat. 12.

3. Attili, S.M., Silva, M.F.M., Nguyen, T., and Ascoli, G.A. (2019). Cell numbers, distribution, shape, and regional variation throughout the murine hippocampal formation from the adult brain Allen Reference Atlas. Brain Struct. Funct. 224, 2883–2897.

4. Ueda, H.R., Dodt, H.-U., Osten, P., Economo, M.N., Chandrashekar, J., and Keller, P.J. (2020). Whole-Brain Profiling of Cells and Circuits in Mammals by Tissue Clearing and Light-Sheet Microscopy. Neuron 106, 369–387.

5. Ragan, T., Kadiri, L.R., Venkataraju, K.U., Bahlmann, K., Sutin, J., Taranda, J., Arganda-Carreras, I., Kim, Y., Seung, H.S., and Osten, P. (2012). Serial two-photon tomography for automated ex vivo mouse brain imaging. Nat. Methods 9, 255–258.

6. Oh, S.W., Harris, J.A., Ng, L., Winslow, B., Cain, N., Mihalas, S., Wang, Q., Lau, C., Kuan, L., Henry, A.M., et al. (2014). A mesoscale connectome of the mouse brain. Nature 508, 207–214.

7. Allen Mouse Brain Connectivity Atlas TECHNICAL WHITE PAPER: OVERVIEW OVERVIEW The Allen Mouse Brain (2016).

8. Wang, Q., Ding, S.-L., Li, Y., Royall, J., Feng, D., Lesnar, P., Graddis, N., Naeemi, M., Facer, B., Ho, A., et al. (2020). The Allen Mouse Brain Common Coordinate Framework: A 3D Reference Atlas. Cell 181, 936-953.e20.

9. Costantini, I., Baria, E., Sorelli, M., Matuschke, F., Giardini, F., Menzel, M., Mazzamuto, G., Silvestri, L., Cicchi, R., Amunts, K., et al. (2021). Autofluorescence enhancement for label-free imaging of myelinated fibers in mammalian brains. Sci. Rep. 11, 8038.

10. Dacosta, R.S., Andersson, H., Cirocco, M., Marcon, N.E., and Wilson, B.C. (2005). Autofluorescence characterisation of isolated whole crypts and primary cultured human epithelial cells from normal, hyperplastic, and adenomatous colonic mucosa. J. Clin. Pathol. 58, 766.

11. Kretschmer, S., Pieper, M., HüTtmann, G., BöLke, T., Wollenberg, B., Marsh, L.M., Garn, H., and KöNig, P. (2016). Autofluorescence multiphoton microscopy for visualization of tissue morphology and cellular dynamics in murine and human airways. Lab. Investig. J. Tech. Methods Pathol. 96, 918–931.

12. Murakami, T.C., Mano, T., Saikawa, S., Horiguchi, S.A., Shigeta, D., Baba, K., Sekiya, H., Shimizu, Y., Tanaka, K.F., Kiyonari, H., et al. (2018). A three-dimensional single-cell-resolution whole-brain atlas using CUBIC-X expansion microscopy and tissue clearing. Nat. Neurosci. 21, 625–637.

13. Neumann, F.R., and Nurse, P. (2007). Nuclear size control in fission yeast. J. Cell Biol. 179, 593–600.

14. Huber, M.D., and Gerace, L. (2007). The size-wise nucleus: nuclear volume control in eukaryotes. J. Cell Biol. 179, 583–584.

15. Seiriki, K., Kasai, A., Hashimoto, T., Schulze, W., Niu, M., Yamaguchi, S., Nakazawa, T., Inoue, K., Uezono, S., Takada, M., et al. (2017). High-Speed and Scalable Whole-Brain Imaging in Rodents and Primates. Neuron 94, 1085-1100.e6.

16. Levy, R.B., Marquarding, T., Reid, A.P., Pun, C.M., Renier, N., and Oviedo, H.V. (2019). Circuit asymmetries underlie functional lateralization in the mouse auditory cortex. Nat. Commun. 10, 2783.

17. Morris, J.A., Jordan, C.L., and Breedlove, S.M. (2008). Sexual dimorphism in neuronal number of the posterodorsal medial amygdala is independent of circulating androgens and regional volume in adult rats. J. Comp. Neurol. 506, 851–859.

18. Morris, J.A., Jordan, C.L., Dugger, B.N., and Breedlove, S.M. (2005). Partial demasculinization of several brain regions in adult male (XY) rats with a dysfunctional androgen receptor gene. J. Comp. Neurol. 487, 217–226.

19. Garcia-Falgueras, A., Pinos, H., Collado, P., Pasaro, E., Fernandez, R., Segovia, S., and Guillamon, A. (2005). The expression of brain sexual dimorphism in artificial selection of rat strains. Brain Res. 1052, 130–138.

20. Herculano-Houzel, S., Ribeiro, P., Campos, L., Silva, A.V. Da, Torres, L.B., Catania, K.C., and Kaas, J.H. (2011). Updated Neuronal Scaling Rules for the Brains of Glires (Rodents/Lagomorphs). Brain. Behav. Evol. 78, 302–314.

21. Herculano-Houzel, S., Catania, K., Manger, P.R., and Kaas, J.H. (2015). Mammalian Brains Are Made of These: A Dataset of the Numbers and Densities of Neuronal and Nonneuronal Cells in the Brain of Glires, Primates, Scandentia, Eulipotyphlans, Afrotherians and Artiodactyls, and Their Relationship with Body Mass. Brain. Behav. Evol. 86, 145–163.

22. Cooke, B.M., Stokas, M.R., and Woolley, C.S. (2007). Morphological sex differences and laterality in the prepubertal medial amygdala. J. Comp. Neurol. 501, 904–915.

23. Qiu, L.R., Fernandes, D.J., Szulc-Lerch, K.U., Dazai, J., Nieman, B.J., Turnbull, D.H., Foster, J.A., Palmert, M.R., and Lerch, J.P. (2018). Mouse MRI shows brain areas relatively larger in males emerge before those larger in females. Nat. Commun. 9, 2615.

24. Lotze, M., Domin, M., Gerlach, F.H., Gaser, C., Lueders, E., Schmidt, C.O., and Neumann, N. (2019). Novel findings from 2,838 Adult Brains on Sex Differences in Gray Matter Brain Volume. Sci. Rep. 9, 1671.

25. Wu, Y., Kirillov, A., Massa, F., Lo, W.-Y., and Girshick, R. (2019). Detectron2.

26. He, K., Gkioxari, G., Dollár, P., and Girshick, R. (2018). Mask R-CNN. ArXiv170306870 Cs.

27. [1512.03385v1] Deep Residual Learning for Image Recognition https://arxiv.org/abs/1512.03385v1.

28. Dutta, A., and Zisserman, A. (2019). The VGG Image Annotator (VIA). CoRR abs/1904.10699.

29. Krizhevsky, A., Sutskever, I., and Hinton, G.E. (2012). ImageNet Classification with Deep Convolutional Neural Networks. In Advances in Neural Information Processing Systems (Curran Associates, Inc.).

30. [1412.6980] Adam: A Method for Stochastic Optimization https://arxiv.org/abs/1412.6980.

31. Jaccard, P. (1912). The Distribution of the Flora in the Alpine Zone.1. New Phytol. 11, 37–50.

32. Amato, S.P., Pan, F., Schwartz, J., and Ragan, T.M. (2016). Whole Brain Imaging with Serial Two-Photon Tomography. Front. Neuroanat. 10.

