## Supplemental Figures for "Sex-, strain and lateral differences in brain cytoarchitecture across a large mouse population"

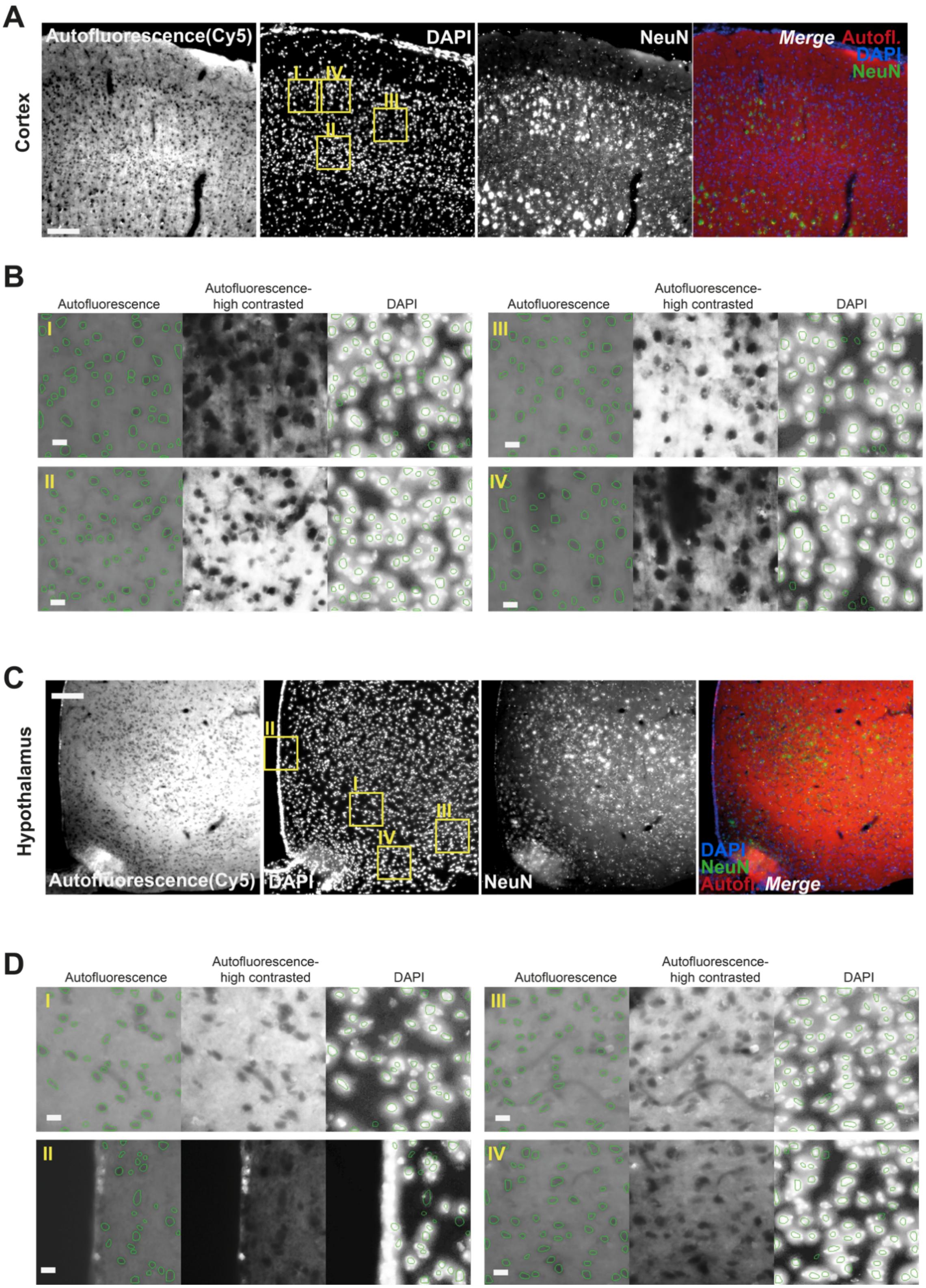


**Supplemental Figure S1 (related to Fig. 1): Autofluorescence signal corresponds to nucleus validation.** A mouse brain was perfused with 4% PFA followed by sectioning, anti-NeuN immunohistochemistry and DAPI counterstaining. Representative image from cortex **A** or hypothalamus **C** showing autofluorescence (Cy5-far red channel), DAPI, NeuN, and the merge of the three channels. **B,D** We applied the same segmentation DNN used for the Allen Mouse Connectivity dataset. Each tile in **B** and **D** shows detected objects on top of the original images (left), autofluorescence high contrast (middle), and DAPI overlayed with the same objects (right). Scale bars, a and c, 50μm; b and d, 10μm.

### Supplemental Figures

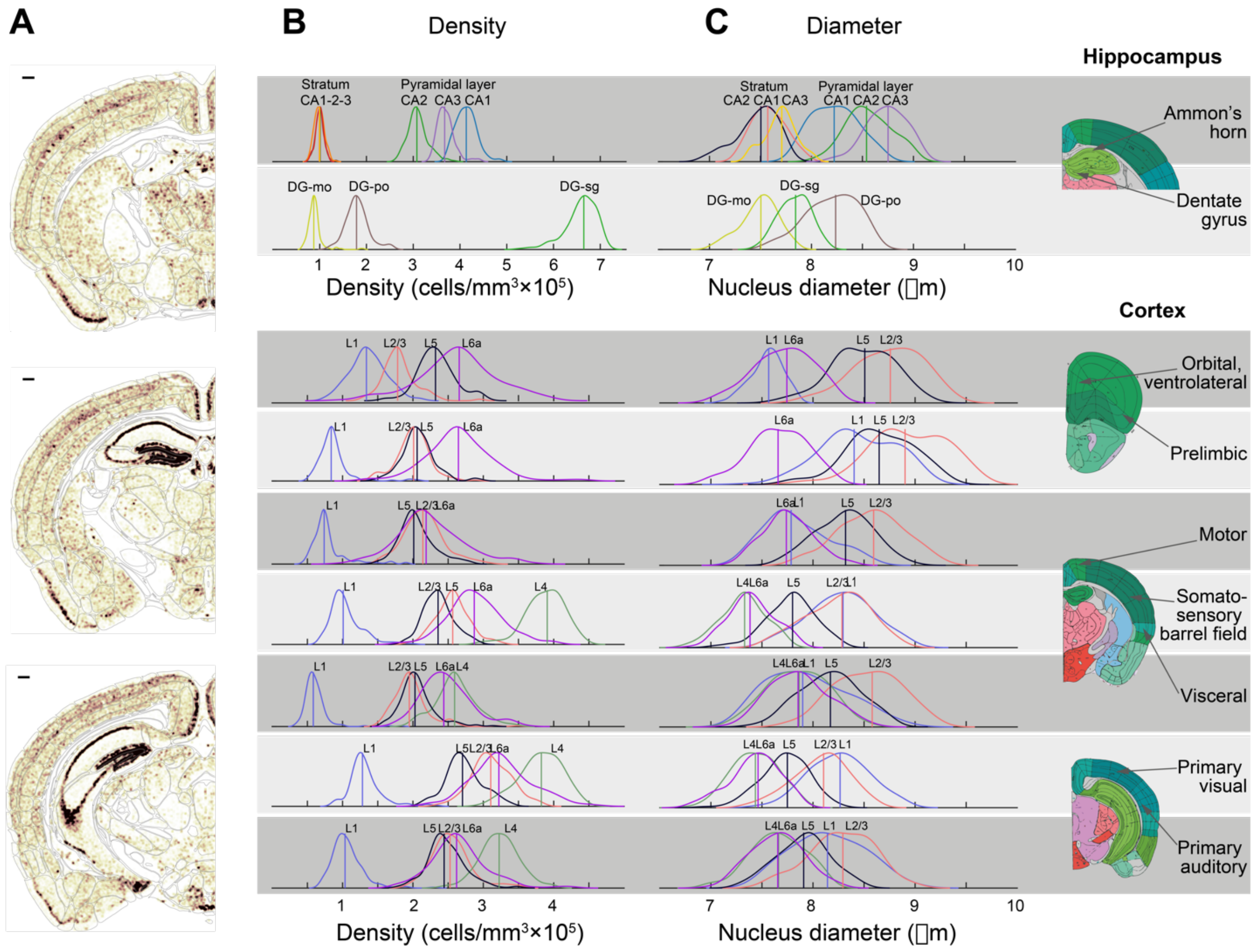


**Supplemental Figure S2 (related to Fig. 1): Density and nucleus diameter along cortical regions. A** Local density is shown as a heat map over the anatomy of three coronal sections of one brain. White, low; dark brown, high local density; scale bars on upper left corners equal 280μm. **B-C** Distribution of cell density **B** and nucleus diameter **C** in the hippocampus and selected cortical regions, in 195 C56BL/6J male mice. The two upper rows show Ammon's horn and the dentate gyrus of the hippocampal formation, and the rows below show examples of cortical regions, each resolved to its cortical layers. On the right, approximate locations of each region are indicated in coronal sections of the AMBA.

### Supplemental Figures

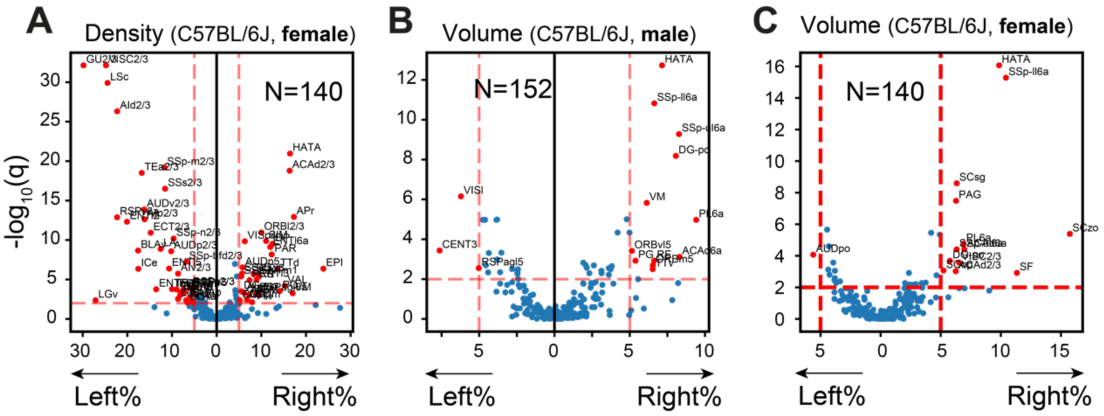


**Supplemental Figure S3 (related to Fig. 2):** Volcano plots showing region-wise comparison of left vs. right hemispheres in C57BL/6: **A** Density comparison in females, and **B-C** volume comparison in males (**B**) and females (**C**).

### Supplemental Figures

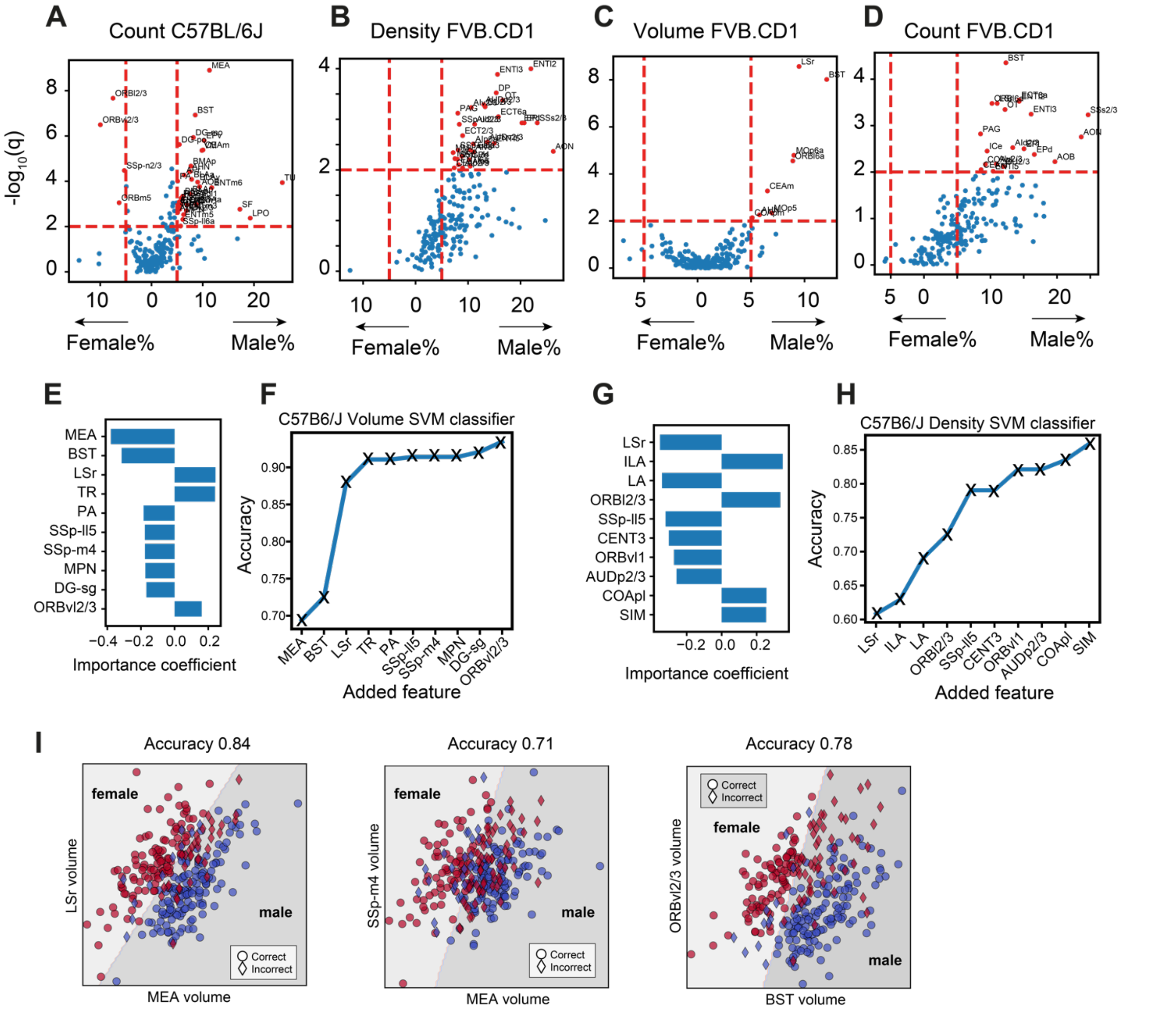

**Supplemental Figure S4** (related to Fig. 3): **A-D** Volcano plots showing per-region statistical testing for male vs. female differences. The horizontal axis represents median differences (%) and the vertical axis displays the q-values (FDR corrected rank-sum p-values by BH procedure in -log10 scale). Red dots correspond to an effect size larger than 5% and q<0.01. **A** Cell count in C57BL/6J, **B** Density in FVB.CD1, **C** Volume in FVB.CD1, **D** Cell count in FVB.CD1. **E** Top 10 regions when constructing a linear SVM volume-based classifier. **F** Accuracy of volume-based classifiers when using the top 10 regions, adding one region at the time. **G-H** same as **E-F,** but for density-based classifiers. **I** Visualization for three examples of two-dimensional SVM based on volume information, as indicated on x and y axes; with accuracies indicated above. The separating line splits the plane into female/male domains. Red/blue markers represent individual males and females, respectively. Circles and diamonds represent correct and incorrect classifications, respectively.

### Supplemental Figures

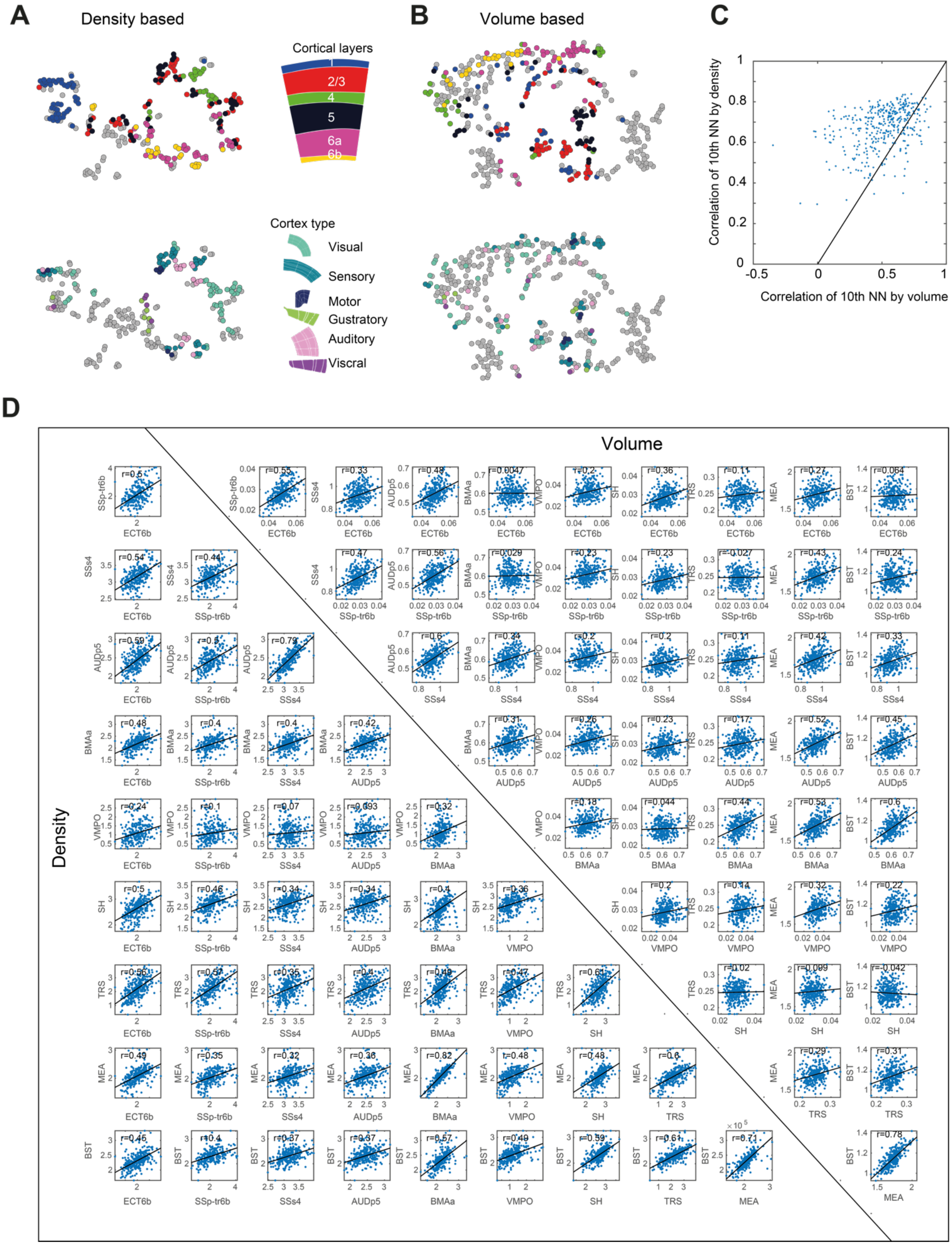


**Supplemental Figure S5 (related to Fig. 5): A** The density based tSNE plot of Fig. 5f color-labelled according to cortical layers (upper) and cortical division (lower). **B** The same for the volume based tSNE plot of Fig. 5g. **C** Correlation to the 10th nearest neighbor for each region when using volume (horizontal axis) or density (vertical axis). **D** Examples for region-region correlations. We show regions ECT6b, SSp-tr6b, SSs4, AUDp5, BMAa, VMPO, SH, TRS, MEA, and BST. Correlations are calculated by density (lower triangle) and volume (upper triangle).
